# D-Retro-Inverso Peptide Candidates for Inhibiting SAA Cardiac Amyloidosis

**DOI:** 10.64898/2026.06.05.730446

**Authors:** Andrew D. Chesney, Lucy M. Coleman, Ulrich H. E. Hansmann

**Affiliations:** Department of Chemistry & Biochemistry, University of Oklahoma, Norman, OK 73019, USA

## Abstract

In a recent study of a mice model it was suggested that after myocardial infarction Serum Amyloid A (SAA) aggregates are formed that contribute to the long-term complications of the infarct, and that a similar mechanism may exist for humans. Motivated by this hypothesis we have designed four peptide candidates that may interfere with formation of SAA3 fibrils, and using all-atom molecular dynamics have evaluated their ability to destabilize SAA fibrils. As the lifetime of peptide drugs can be increased by replacing L-amino acids with their mirror D-amino acids, we have built the peptides from D-amino acids. We identify two of these peptides, DRI-R5S and DRI-H6A, as promising drug candidates.

## I. INTRODUCTION

Following inflammation, Serum amyloid A (SAA) levels increase to upwards of 1000 times normal levels. These high concentrations can lead to amyloidosis, i.e., the misfolding and aggregation of SAA into fibrils deposited into organs such as the liver, kidneys, and the heart. It has been proposed that SAA aggregation may also contribute to the scarring responsible for many of its long-term complications of myocardial infarction, though this has yet to be fully understood.^1–3^ The hypothesis is supported by observation of cardiac SAA amyloidosis post myocardial infarction in a mice model. Here, SAA3 chains^4^ are secreted locally at sites of injury,^5^ and binding to toll-like receptor 2 (TLR2) on macrophages, initiate the enhanced self-production of SAA3 and other cytokines and chemokines.^6, 7^ Saturated lysosomes finally release SAA3 monomers that, enriched with β-sheet motifs prone to aggregation, accumulate into insoluble amyloid protofilaments in the extra cellular matrix.^8^ A similar mechanism may exist in humans as SAAs have high sequence similarities between human and mice, and SAA3 is known to exist in humans as a fusion protein with SAA2.^9^

Assuming that SAA amyloidosis can appear post-myocardial-infarct in humans it becomes important to search for inhibitors of SAA amyloid formation. In previous work we have proposed DRI-R5S as a peptide inhibitor for human SAA aggregation,^10^ and demonstrated computationally its ability to disrupt SAA1 fibrils. The peptide is made of D-amino acids with the reversed sequence of the first five residues of human SAA1, RSFFS, which also is able to disrupt SAA1 fibrils. However, by reversing the target sequence and inverting the chirality, the resulting DRI-peptide is resistant to proteolytic cleavage than RSFFS while maintaining similar sidechain orientations of the native L-enantiomer. Cimini et al., utilized this peptide in their investigations of mouse SAA3 amyloidosis following myocardial infarction. They showed that DRI-R5S decreased the formation of SAA3 amyloid fibrils and improved the cardiac function of the treated mice in comparison to the control group.^4^

The present work complements ongoing experiments in the Cimini and Kishore lab that probe whether DRI-R5S can also dissolve SAA3 fibrils potentially reducing or even reversing post infarct scaring. As there is no experimental structure of SAA3 deposited in the Protein Data Bank (PDB) we have generated the model of a SAA3 fibril shown in **Figure 1** from the homologous human SAA1. This model consists of three layers with two protein chains per layer for a total of six chains. Using virtual screening we then identified and compared a set of hexamer DRI-peptides that could serve as potential inhibitors. For understanding the mechanism by which these peptides block SAA3 aggregation and/or distort SAA3 fibrils, and to identify optimized candidates for future experiments, we studied by molecular dynamics simulations the interaction of these peptides with both single SAA3 chains (**Figure 1a**) the fibril model (**Figure 1b**). To account for potential off-target molecule interactions, we considered interactions with collagen 1a (PDB ID: 2LLP)^11^ and fibronectin (PDB ID: 1FBR)^12^, shown in **Figure 1c** and **Figure 1d**, respectively. Both proteins are present in the heart and important players in the biochemistry of the myocadiac infarction, potentially competing with SAA3 in the interaction with the peptides. Hence, our goal here is not only to evaluate the efficiency of the inhibitor candidates on interfering with SAA aggregation, but also the specificity of that effect.

**Figure 1.**
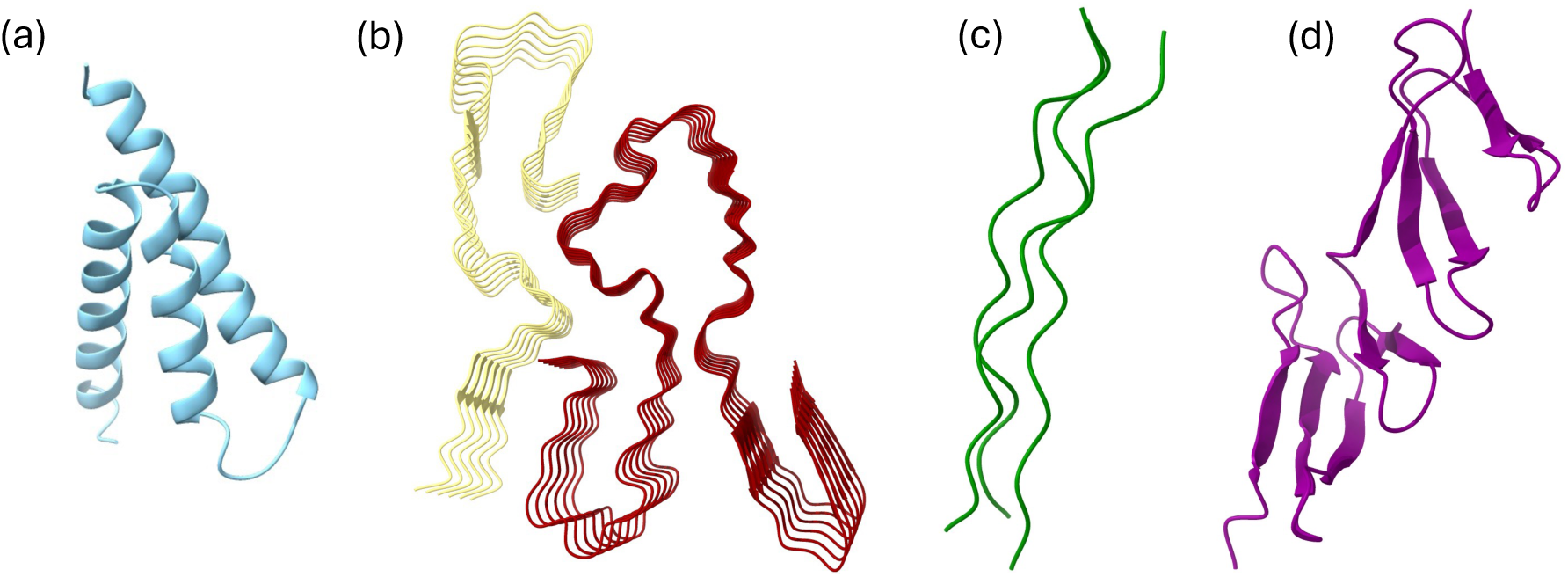
Initial conformations of SAA3 (a) monomer (PDB ID: 6PXZ), (b) fibril model (PDB ID: 7ZKY), (c) collagen 1a (PDB: 2LLP), and (d) fibronectin (PDB: 1FBR). In the SAA3 fibril, protofibril 1 is shown in red and protofibril 2 is shown in light yellow. Structures not shown to scale.

All inhibitor candidates are listed in **Table 1** together with the AutoDock Vina score to target and off-targets. The protocols and techniques used in the set-up and analysis of our simulations are described in the Materials and Methods section that can be found in the Supplemental Information, which also contain the atomic coordinates of the start configurations of all molecular dynamic trajectories. The statistics of all our systems is summarized in **Table S1** of the Supplemental Information.

**Table 1.** AutoDock Vina affinity scores of inhibitor candidates with respect to the target (SAA3 monomer and fibril) and off-target molecules Collagen 1a and Fibronectin.

|  | SAA3 Fibril | Collagen 1a | Fibronectin | SAA Monomer |
| --- | --- | --- | --- | --- |
| DRI-RSFFS (DRI-R5S) | -7.6 | -4.7 | -7.3 | -7.2 |
| DRI-GVWWFF (DRI-G6F) | -9.2 | -4.5 | -6.0 | -7.0 |
| DRI-ERGFFY (DRI-E6F) | -8.7 | -4.3 | -5.6 | -7.1 |
| DRI-HFVWIA (DRI-H6A) | -8.4 | -5.3 | -6.5 | -6.8 |

## II. RESULTS AND DISCUSSION

### Mouse SAA3 Monomer Simulations

We begin our discussion by exploring how the peptides DRI-peptide inhibitor candidates affect the ensemble of SAA3 monomer conformations. Our simulations start from the structure shown in **Figure 1a**, a three-helix bundle. We show in **Figure 2** the average RMSD (Root mean square deviation) in reference to the start configuration as a function of time. This quantity is changing slowly with time, with over the last 100 ns for all DRI-peptides the difference to the start conformation comparable or smaller than in the control, see also **Table 2**. The lowest deviation is seen for DRI-H6A and DRI-R5S.

**Figure 2.**
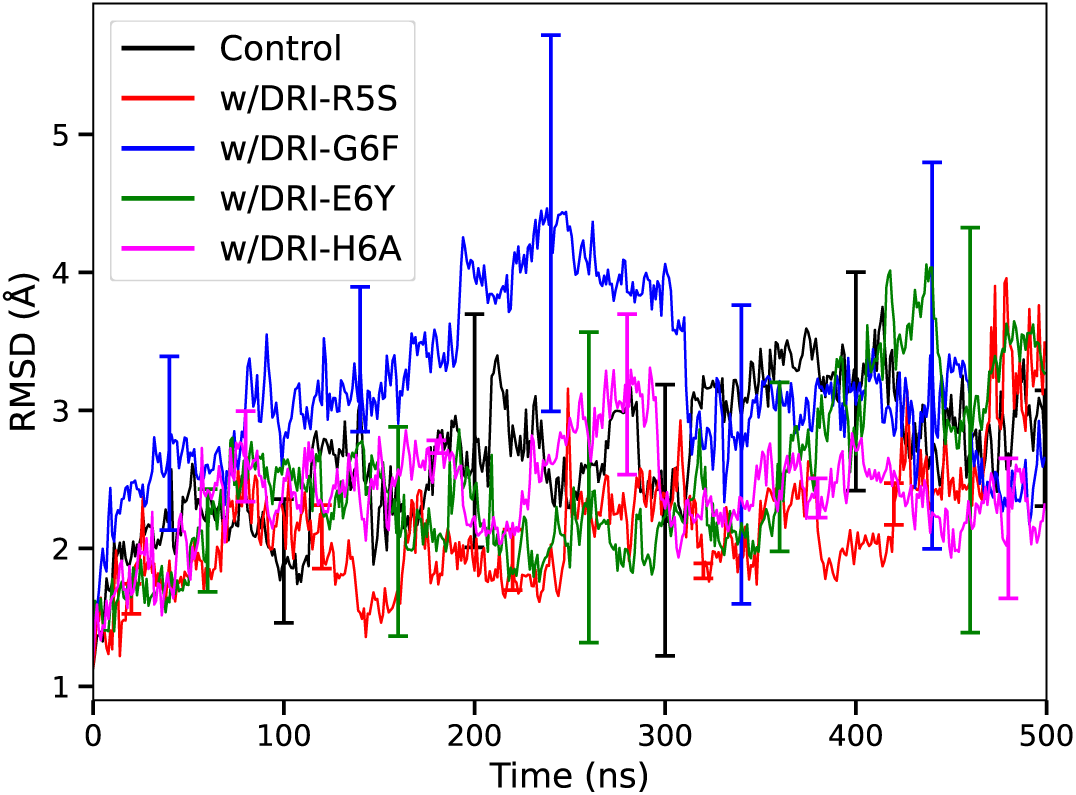
Average backbone RMSD of SAA3 monomer in the absence and presence of inhibitor candidates as function of time, with the bars indicating standard deviation over three trajectories.

**Table 2.** Average Values of RMSD, Rg, Number of Intrastrand, Helical and Non-helical Contacts, and Number of Native Intrastrand, Native Helical, and Native Non-Helical Contacts. Shown are averages over the last 100 ns of three trajectory, with the standard deviation calculated over the three trajectories shown in parentheses.

|  | Control | DRI-R5S | DRI-G6F | DRI-E6Y | DRI-H6A |
| --- | --- | --- | --- | --- | --- |
| RMSD (Å) | 3.0 (5) | 2.7 (1) | 2.9 (7) | 3 (1) | 2.3 (3) |
| Rg (Å) | 12.7 (2) | 12.6 (3) | 12.9 (2) | 12.5 (2) | 13.0 (2) |
| Intrastrand Contacts | 158 (6) | 160 (3) | 156 (4) | 158 (7) | 163 (2) |
| Native Intrastrand Contacts | 135 (6) | 144 (2) | 138 (5) | 140 (10) | 144 (2) |
| Helical Contacts | 113 (2) | 110 (1) | 111 (5) | 106 (5) | 113 (1) |
| Native Helical Contacts | 105 (3) | 104 (2) | 107 (5) | 104 (6) | 109 (1) |
| Non-helical Contacts | 45 (4) | 50 (3) | 45 (3) | 52 (2) | 50 (1) |
| Native Non-helical Contacts | 30 (3) | 39 (1) | 31 (3) | 33 (6) | 35 (1) |

A similar picture is seen in **Figure 3a** for the number of (intra-strand) contacts shown which changes little in presence of the inhibitor candidates but over the last 100 ns drops for the control. Note that we show here only native intra-strand contacts, i.e., such contacts that are seen already in the start conformation and are preserved over the course of the simulation. Consistent with the RMSD plots, we find that such contacts are preserved best for DRI-H6A and DRI-R5S. The intra-strand contacts can be of two types, such between residues close in sequence and characteristic for the three helices seen in the start configuration, and long-range contacts connecting the three helical segments. The loss of intra-strand contacts in the control simulation is mostly from the non-helical contacts, see **Figure 3b** and **3c**. The corresponding averages taken over the last 100 ns are also listed in **Table 2**. While a stabilization of helical contacts (and therefore the three helices) is seen for only DRI-H6A and DRI-G6F, all four inhibitor candidates stabilize non-helical contacts. The effect is largest for DRI-H6A and DRI-R5S and results from the peptides enhancing the interaction of helix 2 (residues 30-45) with both helix 1 (residues 1-26) and helix 3 (residues 47-69), see **Table 3**. These inter-helix interactions are known to be important for the stability of the native structure. Presence of these two candidates also increases the number of contacts between the N-terminus and the rest of the native SAA structure (**Table 3**), stabilizing this segment which unbound otherwise initiates aggregation.

**Figure 3.**
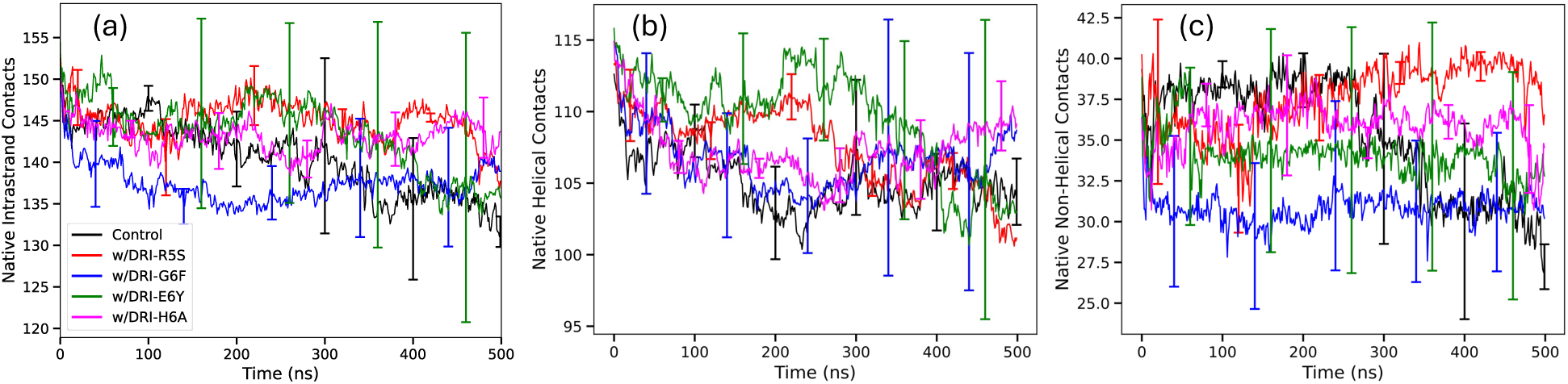
Average number of (a) total, (b) helical, and (c) non-helical contacts between residues in the SAA3 monomer in the absence and presence of DRI peptide inhibitor candidates as function of time, with bars indicating standard deviations over three runs. Counted are only contacts that appear already in the start configuration and are preserved over the trajectory.

**Table 3.**
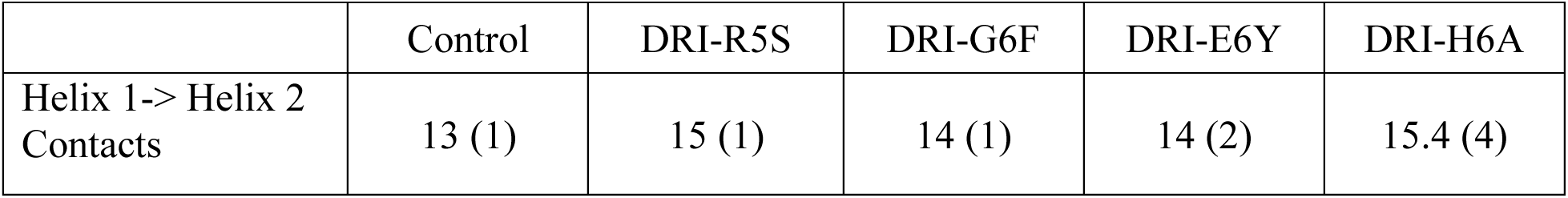
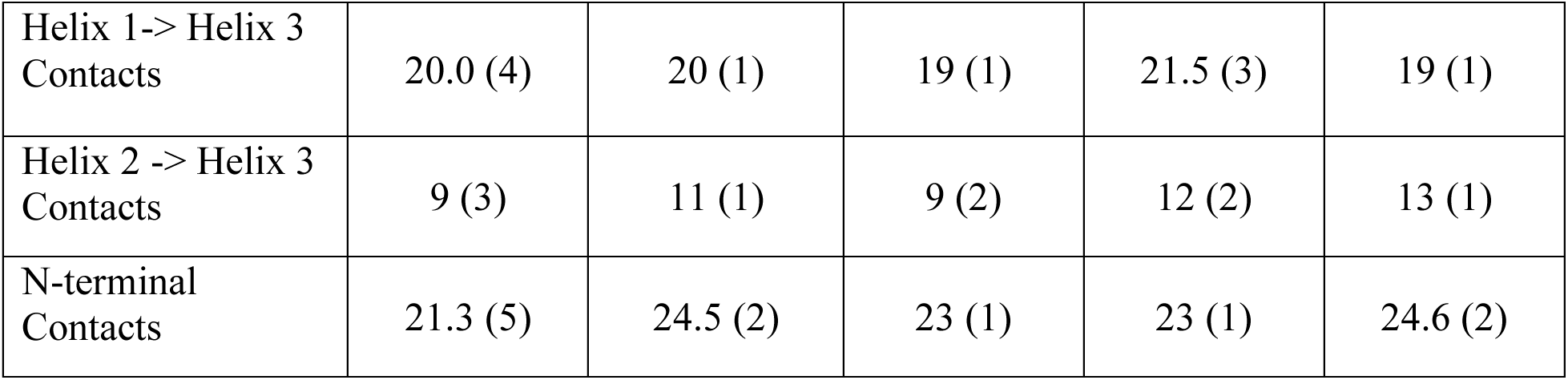
Number of contacts between helices, and between the N-terminal residues (residues 1-11) and the remaining residues of the monomer. Helices are defined as follows: helix 1, residues 1-26; helix 2 residues 30-45; helix 3, residues 47-69. Averages are taken over the last 100 ns, with standard deviations calculated over three independent trajectories shown in parentheses.

|  | Control | DRI-R5S | DRI-G6F | DRI-E6Y | DRI-H6A |
| --- | --- | --- | --- | --- | --- |
| Helix 1-> Helix 2 Contacts | 13 (1) | 15 (1) | 14 (1) | 14 (2) | 15.4 (4) |
| Helix 1-> Helix 3<br>Contacts | 20.0 (4) | 20 (1) | 19 (1) | 21.5 (3) | 19 (1) |
| Helix 2 -> Helix 3<br>Contacts | 9 (3) | 11 (1) | 9 (2) | 12 (2) | 13 (1) |
| N-terminal<br>Contacts | 21.3 (5) | 24.5 (2) | 23 (1) | 23 (1) | 24.6 (2) |

Summarizing our results we find that all four candidates stabilize the native SAA3 monomer conformation of a three-helix-bundle, with DRI-H6A and DRI-R5S both stabilizing contacts between helix 2 and helix 1, or between helix 2 and helix 3, but not between helix 1 and helix 3. In the native structure the N-terminal segment of residues 1-10 is part of helix 1, but when not bound to the rest of the native structure can unfold and misfold into a β-hairpin structure whose presence may trigger aggregation.^13, 14^ Both DRI-H6A and DRI-R5S also support this attachment of the N-terminus therefore providing additional stabilization of the helix bundle. We propose that of the two peptides DRI-H6A is the more efficient inhibitor of SAA3 aggregation as unlike DRI-R5S, it also stabilizes directly the helices (see **Table 2** and **3**) and binds stronger to the monomers, see also **Table 5**.

Previous work^4^ studying the effect of DRI-R5S on SAA aggregation in a mice model provided hints that the peptide not only inhibits formation of SAA fibrils but may also dissolve existing fibrils, a conjecture supported by our earlier computational studies.^10^ For this reason, we evaluate and compare in addition the effect of the four inhibitor candidates on the SAA3 fibril model shown in **Figure 1b**. We note that in **Figure 4a** the average RMSD (root-mean-square deviation) of fibril conformations in reference to the start configuration is larger for the control simulations than for the other systems where inhibitor candidates are present, with the lowest deviation seen in presence of DRI-G6F. The sole exception is DRI-R5S, but here we see a de-stabilizing effect of DRI-R5S only late into the trajectories. We can therefore not exclude the possibility that for the other inhibitors fibrils may be also de-stabilized later in the trajectories. However, such an effect could not come from unfolding of chain conformations. This is because we see in **Figure 4b** no signal for disturbance of the chain geometries in the time evolution of the chain RMSD, which is the average of values calculated separately for all six chains.

**Figure 4.**
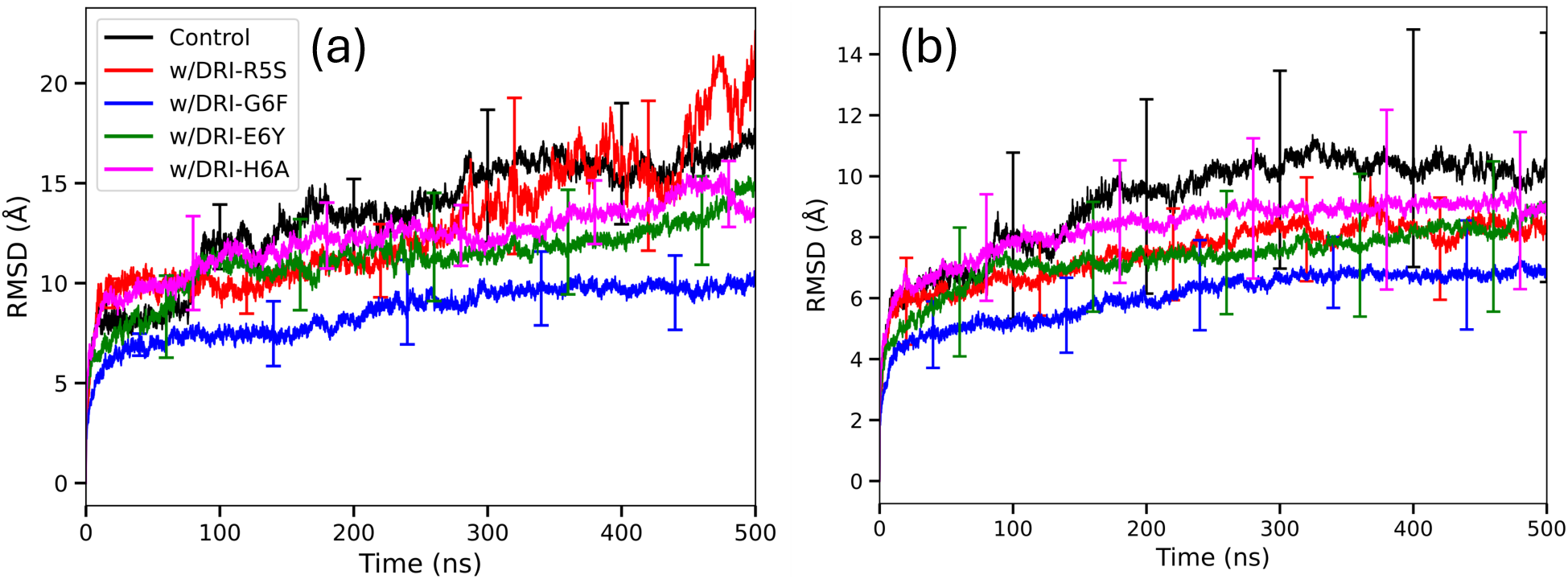
Average (a) global RMSD and (b) chain RMSD of SAA3 backbone atoms in the absence and presence of DRI peptide inhibitor candidates as function of time, with bars indicating standard deviation over three independent runs.

Hence, any de-stabilization of the SAA3 fibril will have to result from disturbance of the arrangement of chains in the SAA3-fibril made by the four inhibitor candidates, i.e., either changing the stacking of chains (in each protofibril) or of their packing (between the two protofibrils) or both. To quantify such potential changes in the arrangement of chains, we have calculated the number of stacking and packing contacts. Stacking contacts are between the layers of the fibril and packing contacts are contacts between the two protofibrils. **Table 4** lists the number of stacking and packing contacts (normalized by dividing through the corresponding numbers at start) averaged over the last 50 ns. The corresponding absolute values are listed in **Table S2**.

**Table 4.** Average normalized final values of total SASA, hydrophobic SASA, hydrophilic SASA, stacking and packing and native stacking and packing contacts for the SAA3 fibril systems. Shown are averages over the last 50 ns of the trajectory, with standard deviations calculated over three independent trajectories shown in parentheses.

| System | Total SASA (Å<sup>2</sup>) | Hydrophobic SASA (Å<sup>2</sup>) | Hydrophilic SASA (Å<sup>2</sup>) | Stacking Contacts | Native Stacking Contacts | Packing Contacts | Native Packing Contacts |
| --- | --- | --- | --- | --- | --- | --- | --- |
| Control | 1.30 (3) | 1.37 (5) | 1.24 (1) | 0.73 (3) | 0.54 (5) | 0.80 (1) | 0.22 (3) |
| DRI-R5S | 1.27 (9) | 1.3 (1) | 1.23 (9) | 0.81 (5) | 0.58 (2) | 0.53 (6) | 0.19 (2) |
| DRI-G6F | 1.26 (5) | 1.19 (7) | 1.33 (3) | 0.80 (5) | 0.49 (4) | 0.42 (8) | 0.06 (3) |
| DRI-E6Y | 1.23 (4) | 1.21 (4) | 1.25 (6) | 0.76 (4) | 0.43 (2) | 0.6 (1) | 0.14 (2) |
| DRI-H6A | 1.27 (5) | 1.25 (8) | 1.29 (3) | 0.76 (3) | 0.43 (1) | 0.5 (3) | 0.19 (5) |

Consistent with our observations for the chain RMSD are the rather small differences in the number of stacking contacts seen between control (73%) and in the presence of inhibitor candidates, varying in number between 76%-81% of the contacts in the initial frame. Note that not all contacts found at the end of the trajectories are the same as the ones seen at start. When considering only native contacts, the corresponding numbers are 43%-58%, and therefore comparable with the one in the control (54%). However, there is a pronounced difference between the control and peptide simulations in the evolution of packing contacts. In the control system is over the final 50 ns the number of packing contacts about 80% of that in the start configuration, and 22% when only native contacts are considered, while in presence of the inhibitor candidates the corresponding numbers vary between 42% to 60% of what is counted in the initial frame, and between 6% and 19% for native contacts. The lowest number of packing contacts is found for DRI-G6F (42%), but visual inspection shows that DRI-G6F is bridging the two protofibrils potentially rather stabilizing the packing of the protofibrils, which may explain the trends observed in RMSD. However, DRI-R5S and DRI-H6A have similarly low numbers (53% and 50%, respectively) agreeing with G6F within the error bars and not acting as bridge between the protofibrils. Note that we see the least reduction in the number of packing contacts for DRI-E6F (60% and 14% for native contacts), which on the other side is with DRI-H6A the peptide causing the least stabilization of stacking contacts.

Hence, for all four inhibitor candidates we see little effect on the stacking of chains but a weakening of the packing of proto fibrils, with the effect strongest for DRI-R5S and DRI-H6A. The effect of this rearrangement of chains can be seen by monitoring how much surface is exposed to the solvent. For this purpose, we list in **Table 4** also the normalized averaged solvent accessible surface area (SASA). Absolute values are given in the supplemental **Table S3.** Folded proteins prioritize burying their hydrophobic residues; and exposed hydrophobic residues increase the chances for aggregation of chains. Hence, we list in **Table 4** not only the total SASA, but also hydrophobic SASA, and hydrophilic SASA. In all cases increases the SASA compared to the initial values, indicating that the final configurations are less compact than the initial structures. When comparing the final normalized hydrophobic SASA, larger difference between the systems are observed. For the control and the DRI-R5S simulations we find an average increase in hydrophobic surface area by about 37% and 30%, respectively, but only by 19% for DRI-G6F which has also the largest increase in the hydrophilic SASA of the SAA3 fibril: by about 33%. Hence, G6F seems to be most effective in interfering with fibril stability. We also note that DRI-G6F has approximately over 80% of its total contacts with the SAA3 fibril remaining while the corresponding numbers for DRI-R5S and DRI-E6Y are approximately 60%, and even less for DRI-H6A with only approximately 50% of its total contacts remaining, see **Table 5** and **S4.** When looking at the evolution of the native, or initial, contacts between the fibril and segments, we note in all simulations a decrease indicating that the peptides are moving away from their initial binding sites. Such a drift is to be expected as there are no restraints applied to the peptides. However, approximately 12% of the initial contacts between DRI-G6F are preserved over the simulation while for the other peptides the frequency of native contacts ranges from 5%-7%. Hence, consisting with the binding energies of **Table 1** we see that DRI-G6F is also interacting stronger with the SAA3 fibril than the other peptides.

**Table 5.** Average normalized number of contacts with peptides calculated for the SAA3 fibril over the last 50ns, and for all other systems over the last 100 ns of the trajectories. Standard deviations are calculated over three independent trajectories and shown in parentheses.

|  |  | DRI-R5S | DRI-G6F | DRI-E6Y | DRI-H6A |
| --- | --- | --- | --- | --- | --- |
| **SAA Monomer** | Contacts | 0.4 (2) | 0.59 (1) | 0.5 (1) | 0.71 (9) |
|  | Native Contacts | 0.09 (7) | 0.15 (3) | 0.10 (8) | 0.2 (1) |
| **SAA Fibril** | Contacts | 0.7 (1) | 0.80 (4) | 0.60 (8) | 0.48 (6) |
|  | Native Contacts | 0.06 (1) | 0.12 (3) | 0.04 (2) | 0.06 (3) |
| **Collagen** | Contacts | 0.11 (4) | 0.25 (7) | 0.3 (1) | 0.11 (3) |
|  | Native Contacts | 0.01 (1) | 0.01 (1) | 0.04 (2) | 0.01 (1) |
| **Fibronectin** | Contacts | 0.2 (1) | 0.61 (3) | 0.2 (1) | 0.4 (1) |
|  | Native Contacts | 0.04 (3) | 0.2 (1) | 0.05 (5) | 0.07 (3) |

#### Simulations of Off Target molecules, Collagen 1a and Fibronectin

To evaluate the potential effects on off-target molecules that were identified by Cimini et. al,^4^ we performed molecular dynamic simulations of collagen 1a (**Figure 1c**) and fibronectin (**Figure 1d**) in the presence and absence of the DRI-peptides. For both molecules we consider the effect on the folded protein structure, studying binding of the peptides to the PDB models (see Methods) and induced unfolding. As for SAA3 monomer and fibril we observe that the peptides are not bound to a specific site but move along the surface of the molecule. The binding contacts between all of the DRI-peptides and collagen 1 decrease even stronger than seen in the case of the SAA3 monomer and fibril, see **Table 5**. This small binding specificity to the initial binding sites, and the neglectable differences in the normalized SASA with the control, see **Table 6** and **S5,** indicate that the effect of the DRI-peptide inhibitor candidates on collagen 1a stability is minimal. For fibronectin is the picture more complex. Binding contacts between fibronectin and DRI-G6F in **Table 5** are much more conserved than these with the other peptides. DRI-G6F has on average ∼60% of the number contacts in the initial frame, while the remaining peptides have less than 20% to 40% of the initial contacts remaining. Hence, DRI-G6F has a higher affinity for fibronectin than the other peptides. However, even for G6F is binding to SAA3 monomer or fibril more favorable. And as for collagen 1 we find again only small differences in SASA values measured in the control and simulations where DRI-peptides are present, see **Table 6** and **S5**. Hence, we conclude that for neither of the four inhibitor candidates are interactions with the two off-target molecules a concern.

**Table 6.** Average normalized final total SASA, hydrophobic SASA, and hydrophilic SASA of collagen 1a and fibronectin in the presence and absence of DRI-Peptides. Standard deviations are calculated over three independent trajectories and shown in parentheses.

| System | Total SASA ($\text{\AA}^2$) | Hydrophobic SASA ($\text{\AA}^2$) | Hydrophilic SASA ($\text{\AA}^2$) |
| --- | --- | --- | --- |
| **Collagen 1a** |  |  |  |
| Control | 1.011 (9) | 0.950 (8) | 1.12 (2) |
| DRI-R5S | 0.96 (5) | 0.95 (5) | 0.96 (8) |
| DRI-G6F | 0.92 (6) | 0.92 (4) | 0.9 (1) |
| DRI-E6Y | 1.0 (2) | 1.0 (2) | 1.1 (2) |
| DRI-H6A | 0.89 (2) | 0.88 (4) | 0.914 (9) |
| **Fibronectin** |  |  |  |
| Control | 0.99 (2) | 1.18 (7) | 0.88 (2) |
| DRI-R5S | 0.98 (2) | 1.10 (2) | 0.90 (3) |
| DRI-G6F | 1.01 (2) | 1.09 (1) | 0.95 (2) |
| DRI-E6Y | 0.98 (3) | 1.08 (4) | 0.91 (3) |
| DRI-H6A | 0.974 (5) | 1.06 (1) | 0.92 (1) |

## III. CONCLUSIONS

Performing all-atom molecular dynamic simulations, we have studied the effect of four small peptides, DRI-R5S, and the newly identified DRI-G6F, DRI-E6Y, DRI-H6A on amyloid formation of mouse SAA3 and the stability of the resulting fibrils. DRI-G6F has a strong binding affinity to both SAA3 monomer and fibril, but also to the off-targets collagen 1a and fibronectin, and as it appears to neither stabilize the SAA3 monomer nor de-stabilize the SAA3 fibril, we conclude that it is not a suitable inhibitor candidate.

The other three peptides have all more favorable binding affinities to SAA3 (both monomer and fibril) than to the off-target collagen 1a and fibronectin. All three candidates stabilize the native SAA3 monomer conformation of a three-helix-bundle by stabilizing non-local contacts between the three helices, with the effect strongest for DRI-H6A and DRI-R5S, therefore reducing the chance for mis-folding and subsequent aggregation. Of the two peptides DRI-H6A may be the more efficient inhibitor of SAA3 aggregation as it stabilizes in addition also the helices themselves and binds stronger to the monomers. On the other hand, the effect of the three peptides on the stability of the SAA3 fibril geometry is more limited, with the peptides interfering more with packing between the protofibril instead of loosening the stacking of the chains. Differences between the three peptides are small, but excluding the somehow mis-leading case of DRI-G6F that bridges the two protofibrils, the loss of packing contacts is lowest for DRI-R5S and DRI-H6A. As DRI-H6A also leads to a larger loss of stacking contacts than the other peptides and to a lower gain in hydrophobic SASA and higher one of hydrophilic SASA than the control, the peptide may not only be more effective than DRI-R5S in inhibiting mis-folding and aggregation of SAA3 monomers, but also in de-stabilizing existing fibrils. Hence, the experimentally observed inhibitor properties of DRI-R5S are likely as much from stabilizing the SAA3 native structure as from de-stabilizing the fibrils and may be shared or even exceeded by DRI-H6A. We therefore suggest using in future experiments not only DRI-R5S as an inhibitor candidate but also DRI-H6A.

## II. MATERIALS AND METHODS

### A. System Preparation

#### Generation of SAA3 Monomer Model

The structure of the SAA3 monomer used in this work was taken from a trimeric structure deposited in the PDB under the ID 6PXZ,^17^ removing residues 90-122. This is because for the similar SAA1 only smaller, cleaved, fragments but not the full peptide form fibrils. The resulting structure is shown in **Figure 1a**.

#### Generation of SAA3 Fibril Model

Presently, there is no fibril structure of mouse SAA3 available from the Protein Data Bank. To generate a structure for use in our simulations, we use the high sequence similarity between human SAA1 and mouse SAA3, to build a model from a vascular variant of human SAA1 with PDB ID 7ZKY,^15^ mutating the residues in this structure to the mouse SAA3 sequence by using UCSF Chimera.^16^ The resulting model is shown in **Figure 1b** and consists of three layers with two protein chains per layer for a total of six chains. Such a system of consisting of two folds and three layers was shown in previous studies to be above the critical threshold required for fibril stability.^10^

#### Generation of Off-Target Molecules Models

In order to evaluate the specificity of our inhibitor candidates we also studied their interactions with two off-target molecules. Start conformations for collagen 1a were generated from the experimentally resolved model for collagen 1a with PDB ID: 2LLP,^11^ and for fibronectin fro. the model with PDB ID: 1FBR.^12^

#### Generation of DRI-Peptides

To identify potential inhibitor candidates we decided to focus on hexapeptides, and given the prohibitively large number of ∼ 20^6 possible combinations of hexapeptides we restricted ourselves to searching the WALTZ-DB 2.0 database^18^ for potential amyloid forming hexapeptides. To generate DRI-peptides, we then reversed the sequences of the resulting 1,402 hexapeptides and generated structure files using PeptideBuilder,^19^ afterwards converting the peptides to their D-enantiomer confirmations by swapping the positions of the β-carbon and α-hydrogen for each residue. The resulting conformations were then energy minimized in gas phase using SANDER from AmberTools.^20^ Binding of the resulting 1,402 DRI-peptides was then screened against the SAA3 fibril model. AutoDock Vina^21^ version 1.1.2 was utilized in a global search as no binding sites of SAA3 are known. A grid search area of 107.309Å x 80.0083Å x 47.3189Å was used to allow ample room for the peptides to move around the SAA3 fibril. To account for potential off-target molecule interactions, the peptides were also screened against collagen 1a (PDB:2LLP) and fibronectin (PDB:1FBR), with search areas of 53.5701Å x 24.4785Å x 26.1194Å and 61.8103Å x 28.7406Å x 37.8707Å, respectively. Following the screening process and guided by the AutoDock Vina affinity scores, the top three inhibitor candidates (high binding affinity to SAA3 monomer and fibril, low binding affinity to the off-targets) were selected for further study. These three peptides selected were DRI-GVWWFF (DRI-G6F), DRI-ERGFFY (DRI-E6Y), and DRI-HFVWIA (DRI-H6A), and added to DRI-RSFFS (DRI-R5S), the peptide inhibitor used in our previous work.^10^ The AutoDock Vina affinity scores of the peptides with respect to SAA3, collagen 1a, and fibronectin are shown in **Table 1**.

The four inhibitor candidates were docked with the SAA3 fibril model in a 1:1 ratio of peptide to chains found in the fibril model using HADDOCK 2.4^22^, resulting in four fibril systems. Similarly, the peptides were docked with SAA3 monomers in a 1:1 ratio, and to explore possible off-target effects with collagen 1a and fibronectin.

### B. General Simulation Protocol

To generate the initial structures for the molecular dynamic simulations performed, the SAA3 fibril and monomer model, collagen 1a (PDB:2LLP), fibronectin (PDB:1FBR), and all the docked structures were converted to GROMACS format using the pdb2gmx module. During this step hydrogen atoms were added to the structures. All chains and peptides were also capped with -NH_3_^+^ and -COO^-^ groups. For the simulations performed the CHARMM 36m-July 2017^23^ all-atom force field was utilized with TIP3P^24^ explicit water molecules. All simulations were performed with GROMACS 2020.4 software package^25^. The structures were then placed in the center of cubic simulation boxes with a minimum distance of 15Å between the solute and the edge of the box. The boxes were then filled with water molecules, and counter ions were added to neutralize the system in accordance with the particle-mesh Ewald method. Na^+^ and Cl^-^ ions were used as the counterions and were set to a physiological concentration of 150 mM NaCl. The total number of atoms and water molecules from each system is listed in **Table S1**. Following the addition of water and counterions, the systems were then energy minimized. The steepest decent method was used, and the systems were minimized for up to 50,000 steps. The systems were then subjected to equilibration at constant volume to set the temperature of the system to 310K for 200ps. Then, at a constant pressure of 1 bar for an additional 200ps. During these equilibration steps the heavy atoms were constrained with a force constant of 1000 kJ/mol•nm^2^.

To main a constant temperature during the simulation a velocity-rescale thermostat^26^ was used with a coupling constant of 0.1 ps. Pressure was maintained by using the Parrinello-Rahman barostat^27^ with a pressure relaxation time of 2 ps. A time step of 2 fs was used for integrating the equations of motion. This was made possible by using the SETTLE^28^ algorithm to insure the water molecules were rigid, as well as restraining protein bonds involving hydrogen atoms to their equilibrium length with the LINCS^29^ algorithm. Periodic boundary conditions were utilized for all systems, as a result the long-range electrostatic interactions were computed with the particle-mesh Ewald method, using a real-space cutoff of 12Å and a 1.6Å Fourier grid spacing. Short-range van der Waals interactions were truncated at 12Å with smoothing beginning at 10 Å. Each system was simulated in triplicate differing by their initial velocity distribution. The length of the trajectories studied for each system is also listed in **Table S1**.

### C. Trajectory Analysis

The molecular dynamics trajectories are analyzed with the GROMACS tools^25^ and MDTraj software^30^. For visualization of trajectory conformations we use VMD^31^ version 1.9.4a57, and UCSF ChimeraX-1.9^32^. Structural evolution of our system was evaluated by measuring the root-mean-square deviation (RMSD), root-mean-square-fluctuation (RMSF), and solvent accessible surface area (SASA), quantities that are calculated using GROMACS tools^25^, for the latter quantity using a spherical probe of 1.4 Å radius. The residue-wise contact frequencies are calculated using GROMACS tools^25^ and MDTraj software^30^, defining contacts by a cutoff 4.5 Å in the closest distance between heavy atoms in a residue pair.

## Supporting information

Start and final configurations

## SUPPORTING INFORMATION

- Five supplemental tables with statistics of the simulations and supporting data
- Start and final configurations of all trajectories in PDB format as text files in a compressed folder

## ACKNOWLEDGMENTS

The simulations in this work were done using the SCHOONER cluster of the University of Oklahoma, XSEDE resources allocated under grant MCB160005 (National Science Foundation).

## Supporting Information

**Table S1.** Simulated Systems.

|  | System | Total Atoms | Water Molecules | Individual Simulation Time (ns) | Total Simulation Time (ns) |
| --- | --- | --- | --- | --- | --- |
| SAA3 | Control | 217,853 | 70,729 | 500 | 1500 |
|  | DRI-R5S | 210,506 | 68,106 |  | 1500 |
|  | DRI-G6F | 209,565 | 67,745 |  | 1500 |
|  | DRI-E6Y | 210,665 | 68,119 |  | 1500 |
|  | DRI-H6A | 211,966 | 68,554 |  | 1500 |
| Collagen 1a | Control | 51,635 | 16,463 | 400 | 1200 |
|  | DRI-R5S | 49,508 | 16,166 |  | 1200 |
|  | DRI-G6F | 49,526 | 16,148 |  | 1200 |
|  | DRI-E6Y | 47,253 | 15,395 |  | 1200 |
|  | DRI-H6A | 50,913 | 16,614 |  | 1200 |
| Fibronectin | Control | 72,606 | 23,689 | 400 | 1200 |
|  | DRI-R5S | 72,195 | 23,522 |  | 1200 |
|  | DRI-G6F | 73,493 | 23,946 |  | 1200 |
|  | DRI-E6Y | 74,025 | 24,124 |  | 1200 |
|  | DRI-H6A | 73,140 | 23,830 |  | 1200 |
| SAA3 Monomer | Control | 53,130 | 17,305 | 500 | 1500 |
|  | DRI-R5S | 53,145 | 17,280 |  | 1500 |
|  | DRI-G6F | 54,326 | 17,665 |  | 1500 |
|  | DRI-E6Y | 54,013 | 17,562 |  | 1500 |
|  | DRI-H6A | 53,913 | 17,529 |  | 1500 |

**Table S2.** Initial and average stacking and packing contacts and native stacking and packing contacts for the SAA3 fibril systems over the last 50 ns. Shown are averages over the last 50 ns of the trajectory, with standard deviations calculated over three independent trajectories shown in parentheses.

|  | Stacking Contacts |  |  | Packing Contacts |  |  |
| --- | --- | --- | --- | --- | --- | --- |
| System | Initial Contacts | Contacts | Native Contacts | Initial Contacts | Contacts | Native Contacts |
| Control | 632 | 502 (20) | 340 (30) | 75 | 35 (4) | 16 (2) |
| DRI-R5S | 614 | 513 (17) | 356 (14) | 76 | 38 (5) | 15 (2) |
| DRI-G6F | 633 | 469 (23) | 312 (23) | 80 | 36 (5) | 5 (2) |
| DRI-E6Y | 640 | 436 (18) | 272 (16) | 73 | 47 (2) | 10 (2) |
| DRI-H6A | 637 | 429 (2) | 274 (3) | 69 | 63 (17) | 13 (3) |

**Table S3.**
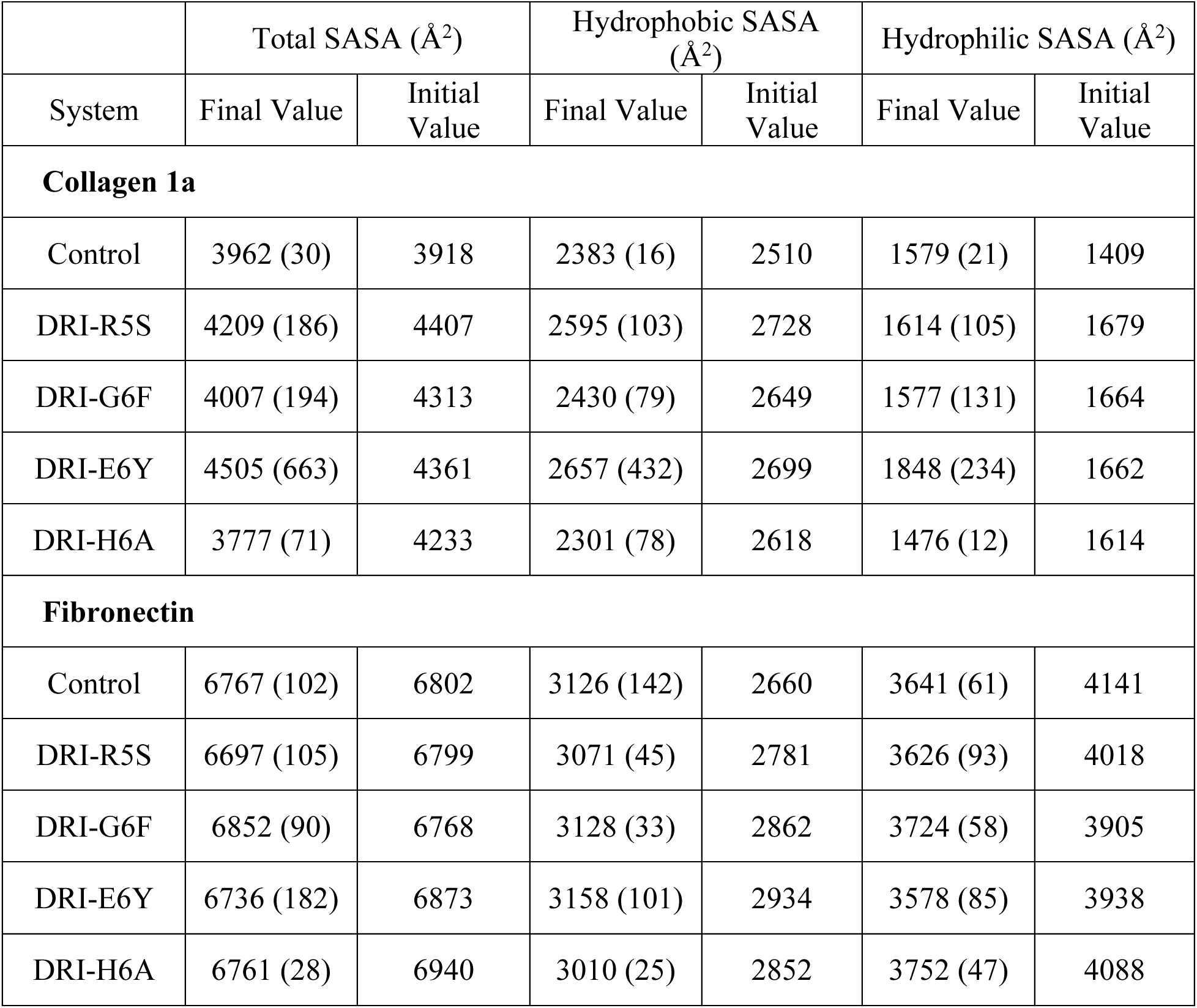
Initial and average final total SASA, hydrophobic SASA, and hydrophilic SASA of collagen 1a and fibronectin in the presence and absence of DRI-Peptides.

|  | Total SASA (Å<sup>2</sup>) |  | Hydrophobic SASA (Å<sup>2</sup>) |  | Hydrophilic SASA (Å<sup>2</sup>) |  |
| --- | --- | --- | --- | --- | --- | --- |
| System | Final Value | Initial Value | Final Value | Initial Value | Final Value | Initial Value |
| **Collagen 1a** |  |  |  |  |  |  |
| Control | 3962 (30) | 3918 | 2383 (16) | 2510 | 1579 (21) | 1409 |
| DRI-R5S | 4209 (186) | 4407 | 2595 (103) | 2728 | 1614 (105) | 1679 |
| DRI-G6F | 4007 (194) | 4313 | 2430 (79) | 2649 | 1577 (131) | 1664 |
| DRI-E6Y | 4505 (663) | 4361 | 2657 (432) | 2699 | 1848 (234) | 1662 |
| DRI-H6A | 3777 (71) | 4233 | 2301 (78) | 2618 | 1476 (12) | 1614 |
| **Fibronectin** |  |  |  |  |  |  |
| Control | 6767 (102) | 6802 | 3126 (142) | 2660 | 3641 (61) | 4141 |
| DRI-R5S | 6697 (105) | 6799 | 3071 (45) | 2781 | 3626 (93) | 4018 |
| DRI-G6F | 6852 (90) | 6768 | 3128 (33) | 2862 | 3724 (58) | 3905 |
| DRI-E6Y | 6736 (182) | 6873 | 3158 (101) | 2934 | 3578 (85) | 3938 |
| DRI-H6A | 6761 (28) | 6940 | 3010 (25) | 2852 | 3752 (47) | 4088 |

**Table S4.** Initial contacts and average number of contacts and native contacts between DRI-peptides and their targets, calculated for the SAA3 fibril over the last 50ns, and for all other systems over the last 100 ns of the trajectories.

|  |  | DRI-R5S | DRI-G6F | DRI-E6Y | DRI-H6A |
| --- | --- | --- | --- | --- | --- |
| **SAA Monomer** | Initial Contacts | 26 | 33 | 33 | 28 |
|  | Contacts | 11 (5) | 19.4 (4) | 16 (4) | 20 (3) |
|  | Native Contacts | 2 (2) | 5 (1) | 3 (3) | 5 (3) |
| **SAA Fibril** | Initial Contacts | 111 | 156 | 166 | 152 |
|  | Contacts | 74 (10) | 124 (6) | 100 (10) | 73 (9) |
|  | Native Contacts | 7 (1) | 19 (4) | 6 (3) | 9 (4) |
| **Collagen** | Initial Contacts | 77 | 65 | 73 | 68 |
|  | Contacts | 8 (3) | 16 (4) | 19 (7) | 8 (2) |
|  | Native Contacts | 0.8 (3) | 0.9 (1) | 3 (2) | 0.9 (4) |
| **Fibronectin** | Initial Contacts | 30 | 28 | 42 | 30 |
|  | Contacts | 6 (4) | 17 (1) | 8 (5) | 11 (3) |
|  | Native Contacts | 1 (1) | 4 (4) | 2 (2) | 2 (1) |

**Table S5.** Initial and average final total SASA, hydrophobic SASA, and hydrophilic SASA of collagen 1a and fibronectin in the presence and absence of DRI-Peptides.

| System | Total SASA (Å<sup>2</sup>) |  | Hydrophobic SASA (Å<sup>2</sup>) |  | Hydrophilic SASA (Å<sup>2</sup>) |  |
| --- | --- | --- | --- | --- | --- | --- |
|  | Initial | Final | Initial | Final | Initial | Final |
| **Collagen 1a** |  |  |  |  |  |  |
| Control | 3918 | 3962 (30) | 2510 | 2383 (16) | 1409 | 1579 (21) |
| DRI-R5S | 4407 | 4209 (185) | 2728 | 2595 (103) | 1679 | 1614 (105) |
| DRI-G6F | 4313 | 4007 (194) | 2649 | 2430 (79) | 1664 | 1577 (131) |
| DRI-E6Y | 4360 | 4505 (663) | 2699 | 2657 (432) | 1662 | 1848 (234) |
| DRI-H6A | 4232 | 3777 (71) | 2618 | 2301 (78) | 1616 | 1476 (12) |
| **Fibronectin** |  |  |  |  |  |  |
| Control | 6802 | 6767 (102) | 2661 | 3126 (142) | 4141 | 3641 (61) |
| DRI-R5S | 6799 | 6697 (105) | 2781 | 3071 (45) | 4018 | 3626 (93) |
| DRI-G6F | 6768 | 6852 (90) | 2862 | 3128 (33) | 3905 | 3724 (58) |
| DRI-E6Y | 6873 | 6736 (182) | 2934 | 3158 (101) | 3938 | 3578 (85) |
| DRI-H6A | 6940 | 6761 (28) | 2853 | 3010 (25) | 4088 | 3752 (47) |

