## Supplementary material for "D-Retro-Inverso Peptide Candidates for Inhibiting SAA Cardiac Amyloidosis": Start and final configurations: README.txt

This directory contains (besides this README.txt) four folders (Fibril, Monomer, Collagen, Fibronectin) that contain files with the atomic coordinates (in PDB format) of aS structures discussed in the article.

Fibril;
This folder contains five sub-directories containing all SAA3 fibril configurations:

1) control:
Atomic coordinates (in PDB format) of the start and final configurations of the control SAA3 fibril simulations:
saa3_f_control_1_initial : Initial structure of SAA3 fibril replica 1
saa3_f_control_2_initial : Initial structure of SAA3 fibril replica 2
saa3_f_control_3_initial : Initial structure of SAA3 fibril replica 3
saa3_f_control_1_final   : Final structure of SAA3 fibril replica 1
saa3_f_control_2_final   : Final structure of SAA3 fibril replica 2
saa3_f_control_3_final   : Final structure of SAA3 fibril replica 3

2) R5S:
Atomic coordinates (in PDB format) of the start and final configurations of the SAA3 fibril simulations with R5S:
saa3_f_R5S_1_initial : Initial structure of SAA3 fibril with R5S replica 1
saa3_f_R5S_2_initial : Initial structure of SAA3 fibril with R5S replica 2
saa3_f_R5S_3_initial : Initial structure of SAA3 fibril with R5S replica 3
saa3_f_R5S_1_final   : Final structure of SAA3 fibril with R5S replica 1
saa3_f_R5S_2_final   : Final structure of SAA3 fibril with R5S replica 2
saa3_f_R5S_3_final   : Final structure of SAA3 fibril with R5S replica 3

3) G6F:
Atomic coordinates (in PDB format) of the start and final configurations of the SAA3 fibril simulations with G6F:
saa3_f_G6F_1_initial : Initial structure of SAA3 fibril with G6F replica 1
saa3_f_G6F_2_initial : Initial structure of SAA3 fibril with G6F replica 2
saa3_f_G6F_3_initial : Initial structure of SAA3 fibril with G6F replica 3
saa3_f_G6F_1_final   : Final structure of SAA3 fibril with G6F replica 1
saa3_f_G6F_2_final   : Final structure of SAA3 fibril with G6F replica 2
saa3_f_G6F_3_final   : Final structure of SAA3 fibril with G6F replica 3

4) E6Y:
Atomic coordinates (in PDB format) of the start and final configurations of the SAA3 fibril simulations with E6Y:
saa3_f_E6Y_1_initial : Initial structure of SAA3 fibril with E6Y replica 1
saa3_f_E6Y_2_initial : Initial structure of SAA3 fibril with E6Y replica 2
saa3_f_E6Y_3_initial : Initial structure of SAA3 fibril with E6Y replica 3
saa3_f_E6Y_1_final   : Final structure of SAA3 fibril with E6Y replica 1
saa3_f_E6Y_2_final   : Final structure of SAA3 fibril with E6Y replica 2
saa3_f_E6Y_3_final   : Final structure of SAA3 fibril with E6Y replica 3

5) H6A:
Atomic coordinates (in PDB format) of the start and final configurations of the SAA3 fibril simulations with H6A:
saa3_f_H6A_1_initial : Initial structure of SAA3 fibril with H6A replica 1
saa3_f_H6A_2_initial : Initial structure of SAA3 fibril with H6A replica 2
saa3_f_H6A_3_initial : Initial structure of SAA3 fibril with H6A replica 3
